# CD177 is a novel IgG Fc receptor and CD177 genetic variants affect IgG-mediated function

**DOI:** 10.1101/2024.01.07.574546

**Authors:** Yunfang Li, Jianming Wu

## Abstract

CD177 plays an important role in the proliferation and differentiation of myeloid lineage cells including neutrophils, myelocytes, promyelocytes, megakaryocytes, and early erythroblasts in bone marrow. CD177 deficiency is a common phenotype in humans. Our previous studies revealed genetic mechanisms of human CD177 deficiency and expression variations. Up to now, immune functions of CD177 remain undefined. In the current study, we revealed human IgG as a ligand for CD177 by using flow cytometry, bead-rosette formation, and surface plasmon resonance (SPR) assays. In addition, we show that CD177 variants affect the binding capacity of CD177 for human IgG. Furthermore, we showed that the CD177 genetic variants significantly affect antibody-dependent cell-mediated cytotoxicity (ADCC) function. The demonstration of CD177 as a functional IgG Fc-receptor may provide new insights into CD177 immune function and genetic mechanism underlying CD177 as biomarkers for human diseases.

## Introduction

IgG Fc receptors (or FcγRs) are surface receptors of immune cells to recognize IgG immune complexes and antibody-opsonized targets. FcγRs are essential for antibody-dependent cell-mediated phagocytosis (ADCP) and antibody-dependent cellular cytotoxicity (ADCC) to eradicate pathogens and infected/mutated target cells. The interaction between FcγRs on immune cells and target-bound antibodies initiates signal cascades for ADCC and ADCP (1). Humans possess a variety of FcγRs to regulate immune responses (2). Genetic polymorphisms of human FcγR genes significantly affect immune responses and disease susceptibility (3–6).

CD177 (NB1 and PRV-1) is a GPI-linked cell surface protein (7, 8) on bone marrow cells (9, 10). CD177 plays an important role in the proliferation and differentiation of myeloid lineage cells including neutrophils, myelocytes, promyelocytes, megakaryocytes, and early erythroblasts (9, 11–14). In peripheral blood, CD177 is exclusively expressed on neutrophils that serve as the first line of defense against infections and the primary mediator for inflammation. CD177 is a biomarker for a number of human diseases. CD177 expression is the most dysregulated in sepsis patients (15), indicating a role in infection. CD177^+^ neutrophil play a protective role in the development of inflammatory bowel disease (16), indicating a role of CD177 in inflammation. Low percentages of CD177^+^ neutrophils are significantly associated with myelodysplastic syndrome and chronic myelogenous leukemia (10, 17). As an important biomarker for polycythemia vera and essential thrombocythemia (14, 18, 19), CD177 plays a pivotal role in myeloid proliferation and differentiation (11–13). CD177 overexpression was significantly correlated with a favorable prognosis for gastric cancer (20) and CD177^+^ neutrophils suppress epithelial cell tumourigenesis in colitis-associated cancer and predict good prognosis in colorectal cancer (21), pointing to a protective role of CD177 in cancer. Up to now, the mechanism with which CD177 influences tumor outcome is unknown (20).

In the previous studies, we delineated genetic mechanisms of CD177 deficiency and expression variations (22, 23). CD177 mediates neutrophil activation triggered by autoantibodies (24), but the physiological ligands and immune functions of CD177 remain incompletely understood. In the current study, we revealed that CD177 is a functional FcγR and that CD177 genetic variants affect CD177 immune function. Our study may provide novel insights into CD177-mediated immune functions and the mechanism underlying the roles of CD177 as a disease biomarker for human diseases.

## Materials and Methods

### Reagents

Human serum IgG (hIgG) and IgA (hIgA) were purchased from Sigma Chemical Co. (St. Louis, MO). FITC (or APC)-conjugated anti-CD177 mAb (clone MEM-166, mouse IgG1) and respective isotype controls were from BioLegend (San Diego, CA). FITC-conjugated goat anti-human Ig(H+L) F(ab’)_2_ were from Jackson ImmunoResearch (West Grove, PA). Human IgG1κ, IgG2κ, IgG3κ, and IgG4κ (purity >95%) were obtained from Athens Research and Technology (Athens, GA). Human mAb trastuzumab/Herceptin manufactured by Genentech (South San Francisco, CA) was obtained through the University of Minnesota Boynton Pharmacy as previously described (25). This study also used puromycin from InvivoGen (San Diego, CA), geneticin (G418) from Life Technologies (Grand Island, NY), and QuikChange Site-directed mutagenesis kit Stratagene (La Jolla, CA).

### Generation of CD177 expression constructs

The human CD177 expression constructs were generated by cloning *Hind* III/*Xba* I-flanked full-length *CD177* cDNA into the eukaryotic expression vector pcDNA3 (Gibco BRL) as previously described (22). For the generation of CD177 retroviral expression constructs, CD177 cDNA inserts from CD177-pcDNA3 expression constructs were amplified by PCR using the forward primer (5’-CTC TAG ACT GCC <u>GGA TCC</u> ACC AGC CAC AGA CGG GTC ATG AG-3’) and the reverse primer (5’-CGA GGC CTG CAG <u>GAA TTC</u> GAG GTC AGA GGG AGG TTG AGT GTG-3’) (the underlined nucleotides indicate *Bam* HI and *EcoR* I restriction sites, respectively). The full-length CD177 cDNA inserts were sub-cloned into pBMN-I-EGFP retroviral vector digested with *Bam* HI and *EcoR* I restriction enzymes using InFusion HD Cloning Kit (Takara Bio USA, Mountain View, Ca) according to the product manual. For the production of soluble CD177, CD177 cDNA coding for amino acids 1-408 was amplified by PCR from the CD177-pcDNA3 expression construct using the forward primer 5’-GAG <u>TCT AGA</u> ACC AGC CAC AGA CGG GTC ATG AG −3’ (the underlined nucleotides indicate an *Xba* I restriction sites) and the reverse primer 5’-CCG <u>GGA TCC</u> TTA ATG ATG ATG ATG ATG ATG ACC TCC CTC ATG CTG AGA GGC AG-3’ (the underlined nucleotides indicate a *Bam* HI restriction sites). The amplified CD177 cDNA fragment with the addition of 6×histidine-tag at the carboxyl terminus was digested by *Xba* I and *Bam* HI before being cloned into a pLenti-3F vector as previously described (26). Nucleotide sequences of all CD177 expression constructs were confirmed by direct Sanger sequencing on an ABI 3730xl DNA Analyzer with BigDye v3.1 Sequencing kit (Applied Biosystems, Inc., Foster City, CA).

### Generation of 293 cell lines expressing CD177

Human embryonic kidney cell line 293 cells (ATCC#CRL-1573, Manassas, VA) were cultured with the DMEM medium containing 10% fetal calf serum, L-glutamine (2 mM), and 1× Pen/Strep (Life Technologies) in 5% CO_2_ incubator at 37 °C. Transfection reactions were carried out in 100-mm cell culture dishes with the plasmid DNA (20 μg) purified with OMEGA Plasmid Maxi Kit (Omega Bio-Tek, Norcross, GA) and 40 μl of Lipofectamine 2000 reagent (Life Technologies). Transfected cells were cultured in DMEM medium for two days before the supplement of G418 (final concentration: 1 mg/ml) for the selection of stable cell lines. The polyclonal cells surviving the G418 selection were sorted with StemCelll EasySep Cell Sorter and the expression of CD177 on the transfected 293 cell lines was determined with FITC-conjugated anti-CD177 mAb as described previously (22).

### Generation of NK-92 cell lines expressing CD177

For retrovirus production, PhoenixA cells were cultured in DMEM supplemented with 10% FBS to ∼ 80% confluence. Transfections of PhoenixA cells were carried out in 100-mm cell culture dishes with pBMN-I-EGFP construct plasmid DNA (20 μg) and 40 μl of Lipofectamine 2000 reagent (Life Technologies). Transfected cells were cultured in DMEM medium for two days before culture supernatants containing retroviral particles were harvested. Cell culture supernatants containing pseudo retrovirus particles were filtered through 0.4 µm syringe filter and used for transduction. IgG Fc receptor-deficient NK-92 (ATCC# CRL-2407, Manassas, VA) cells were maintained in Alpha Minimum Essential medium supplemented with 12.5% heat-inactivated fetal bovine serum (FBS), 12.5% heat-inactivated horse serum, 1× Antibiotic-Antimycotic, and 100 U/ml IL-2 (R&D Systems, Minneapolis, MN). NK-92 cells were passaged in 2–3 day interval to maintain the cell density at 2.5 × 10^5^ – 1 × 10^6^ cells/ml. NK-92 cell transduction was carried out as previously described (27). GFP^+^ NK-92 cells were sorted with FACSAria II cell sorter (BD Biosciences).

### Flow cytometry

The CD177 expression on cells were determined with flow cytometry analysis as described (22). Cells stained with anti-CD177 mAb or isotype control were analyzed on a FACSCelesta flow cytometer (BD Biosciences). The FlowJo software (Tree Star Inc.) was used to evaluate flow cytometry data.

### IgG binding assay

The binding of human IgG (hIgG) to 293 cells expressing CD177 was measured by quantitative flow cytometry as previously described (28, 29). 293 cells were washed twice with cold PBS containing 1% BSA (wash buffer) and re-suspended at the density of 2×10^6^ /ml in wash buffer. An aliquot of 100 µl (2×10^5^) of 293 cells was transferred to a flow tube and centrifuged at 400 ×g for 6 min to pellet cells. After removal of supernatant, 100 µl of hIgG or hIgA diluted at various concentrations in wash buffer was added to each tube. The cells were vortexed and incubated on ice for 30 minutes. Cells were then washed three times with wash buffer and were stained with 100 µl of FITC-conjugated goat anti-human IgG (H+L) antibody (1 μg/ml final concentration) on ice for 30 min. The cells were washed and fixed with 1% formaldehyde before being analyzed on a Celesta flow cytometer (BD Biosciences). For phosphatidylinositol-specific phospholipase C (PI-PLC) treatment, 293 cells (10^6^ cells) expressing CD177 were treated by 0.2 unit PI-PLC (cat# P-6466, ThermoFisher Scientific, Waltham, Massachusetts) in 0.5 ml PBS for 1 h at 37°C. PI-PLC treated cells were washed twice with wash buffer for the IgG binding assay at the final IgG concentration of 100 µg/ml. For mAb blockade analysis, 293 cells expressing CD177 were incubated with anti-CD177 mAb MEM-166 (10 ug/ml final concentration) for 30 min at room temperature. MEM-166 mAb treated cells were washed twice with wash buffer before being used for the IgG binding assay at the final IgG concentration of 200 µg/ml.

### Production and purification of soluble CD177 (sCD177)

Pseudo-lentiviral particles containing sCD177 construct were generated using 293T cells and packaging vectors pMD2.G and pCMV-dR8.74psPAX2 (Addgene). Pseudo-lentiviral particles were transduced into the FreeStyle 293-F cells (ThermoFisher Scientific). The stable 293-F cell line was established under 1.0 µg/ml puromycin selection. The 293-F cells stably expressing sCD177 were cultured in the FreeStyle™ 293 Expression Medium (ThermoFisher Scientific) and cell culture supernatant was harvested when cell density reached 2.5×10^6^/ml. sCD177 was purified from cell culture supernatants using a two-step purification procedure including Ni-affinity chromatography and Fast protein liquid chromatography. For the first step, His-tagged proteins in cell culture supernatant was purified on a HisTrap HP His tag protein purification column (Cytiva, Marlborough, MA) according to the manufacture’s protocol. sCD177 from the first step purification was injected into a Superdex 200 Increase 10/300 GL column (Millipore-Sigma) on an AKTA pure protein purification system (Cytiva) for the second step purification. The purity of proteins was >95% as determined by SDS-PAGE (Supplemental Figure 1).

### Surface plasmon resonance (SPR) analyses

A Biacore S200 instrument (GE Healthcare) was used to determine interaction affinity between sCD177 with human IgG subclasses (IgG1, IgG2, IgG3, and IgG4). In SPR analyses, pure CD177 protein was dialyzed against 1×HBS-P+ buffer (cat# BR100671, Cytiva, Uppsala, Sweden) as previously described (30). An N-hydroxysuccinimide ester was formed on a CM5 sensor chip (GE Healthcare) surface according to a procedure recommended by the manufacturer. Pure human IgG proteins diluted in 100 mM Sodium Acetate (pH 4.5) were immobilized at acidic pH, resulting in the following densities: IgG1: 254 RU, IgG2: 259 RU, IgG3: 185 RU, and IgG4: 239 RU on a CM5 chip. In multicycle SPR kinetic analysis, sCD177 at concentrations of 125, 250, 500, 1,000, 2,000, 4,000, 8,000, and 16,000 nM in 1×HBS-P+ buffer were injected into flow cells at a flow rate of 30 µl/min, with a contact and dissociation time of 250 and 600 seconds, respectively. After each assay cycle, the sensor chip surface was regenerated by 10 mM NaOH. Binding response was recorded as resonance units continuously, with background binding automatically subtracted. The kinetic constants (kon, koff, t1/2) were determined.

Association (kon) and dissociation (koff) rate constants were calculated by a global fitting analysis assuming a Langmuir binding model and a stoichiometry of (1:1). The kinetic dissociation constant (K_D_) was determined from the ratio of the rate constants (K_D_ = koff/kon). K_A_ was calculated by studying the concentration-dependence of the steady-state signal reached at the end of the injection using BIA evaluation software (GE Healthcare).

### Human IgG-coated beads-rosette formation assay

Human IgG (hIgG) was coupled to M-450 Epoxy Dynabeads (cat# 14011, Thermo Fisher Scientific) according to the product manual. Briefly, 1 ml (4×10^8^) Dynabeads were transferred to a microfuge tube and washed twice with 1 ml PBS (pH 7.4) on a magnet. The washed beads were mixed with 200 µg of hIgG in 1 ml of PBS and incubated at room temperature for 15 min. Following the incubation, 20 µl of 5% BSA (bovine serum albumin) was added to the mixture to the final concentration of 0.1% BSA. The beads-hIgG-BSA mixture was continuously incubated for 24 hours at room temperature with rotations on a vertical rotator. After the completion of coupling process, beads were placed on a magnet for the removal of supernatant containing unbound hIgG and BSA. hIgG-coupled beads were washed three times on a magnet with 1 ml of pH 7.4 PBS containing 0.1% BSA and 2 mM EDTA. The hIgG-coupled beads was re-suspended in 1 ml of pH 7.4 PBS containing 0.1% BSA and 2 mM EDTA to obtain 4×10^8^ beads/ml. The rosette formation assay was performed as described (31). IgG-coated beads were mixed with 293 cells at a ratio of 3.5 (3.5 beads per cell). All cells with the attachment of two or more beads were counted in the beads-rosette assay to determine the binding capacity of CD177 alleles.

### ADCC assay

DELFIA EuTDA Cytotoxicity Reagents kit (PerkinElmer, Waltham, MA) was used for ADCC assay as previously described (25, 32). Briefly, SKOV-3 target cells were labeled with Bis(acetoxymethyl)-2-2:6,2 terpyridine 6,6 dicarboxylate (BATDA) for 30 min in their culture medium before being extensively washed in 10 ml DMEM culture medium for three times. The BATDA-labeled cells were transferred into a 96-well non-tissue culture-treated U-bottom plates at a density of 8 × 10^3^ cells/well.

Trastuzumab was added directly to the SKOV-3 cells at the final concentration of 5 μg/ml and NK-92 cells were added at the E:T ratio of 20:1. Effector and target cells in each well were mixed by pipetting up-and-down five times. The plates were then centrifuged at 400 × g for 1 min and subsequently incubated for 2 h in a humidified 5% CO_2_ atmosphere at 37°C. At the end of the incubation, the plates were centrifuged at 500 × g for 5 min and 20 µl of supernatant from each well was transferred to a 96 well DELFIA Yellow Plate (PerkinElmer) and combined with 200 µl of europium. Fluorescence was measured by time-resolved fluorometry using a BMG Labtech CLARIOstar plate reader (Cary, NC).

BATDA-labeled target cells alone with or without therapeutic antibodies were cultured in parallel to assess spontaneous lysis and in the presence of 1% Triton-X to measure maximum lysis. ADCC for each sample is represented as % specific release and was calculated using the following formula: Percent Specific Release = (Experimental release – Spontaneous release)/(Maximal release – Spontaneous release)*100.

### Statistical analysis

ANOVA was used to analyze binding capacity and ADCC differences of CD177 alleles.

## Results

### Human IgG specifically binds to cells expressing CD177

To establish CD177 as a bona fide IgG Fc receptor, we carried out flow cytometry-based IgG binding assay using 293 cells expressing CD177 and vector control. As shown in **Fig. 1**, cells transfected with vector control do not bind human IgG (**Fig. 1B**). In addition, human IgA does not bind to cells expressing CD177 (**Fig. 1C**). However, cells expressing CD177 bound human IgG in a dose-dependent fashion (**Fig. 1D**), indicating that CD177 is an IgG receptor. CD177 is a GPI-linked cell surface glycoprotein and PI-PLC could cleave CD177 from the cell surface (8). As shown in **Fig. 2A**, PI-PLC treatment dramatically reduced CD177 intensity on cell surface. Importantly, cells treated with PI-PLC almost lost the binding ability to human IgG (**Fig. 2B**) compared to the same cell without PI-PLC treatment. In addition, treatment of cells with anti-CD177 monoclonal antibody (mAb MEM-166) significantly inhibited the binding of human IgG to the cells expressing CD177 (**Fig. 2C**). Taken together, our data confirmed the interaction between CD177 and human IgG.

**Fig. 1.**
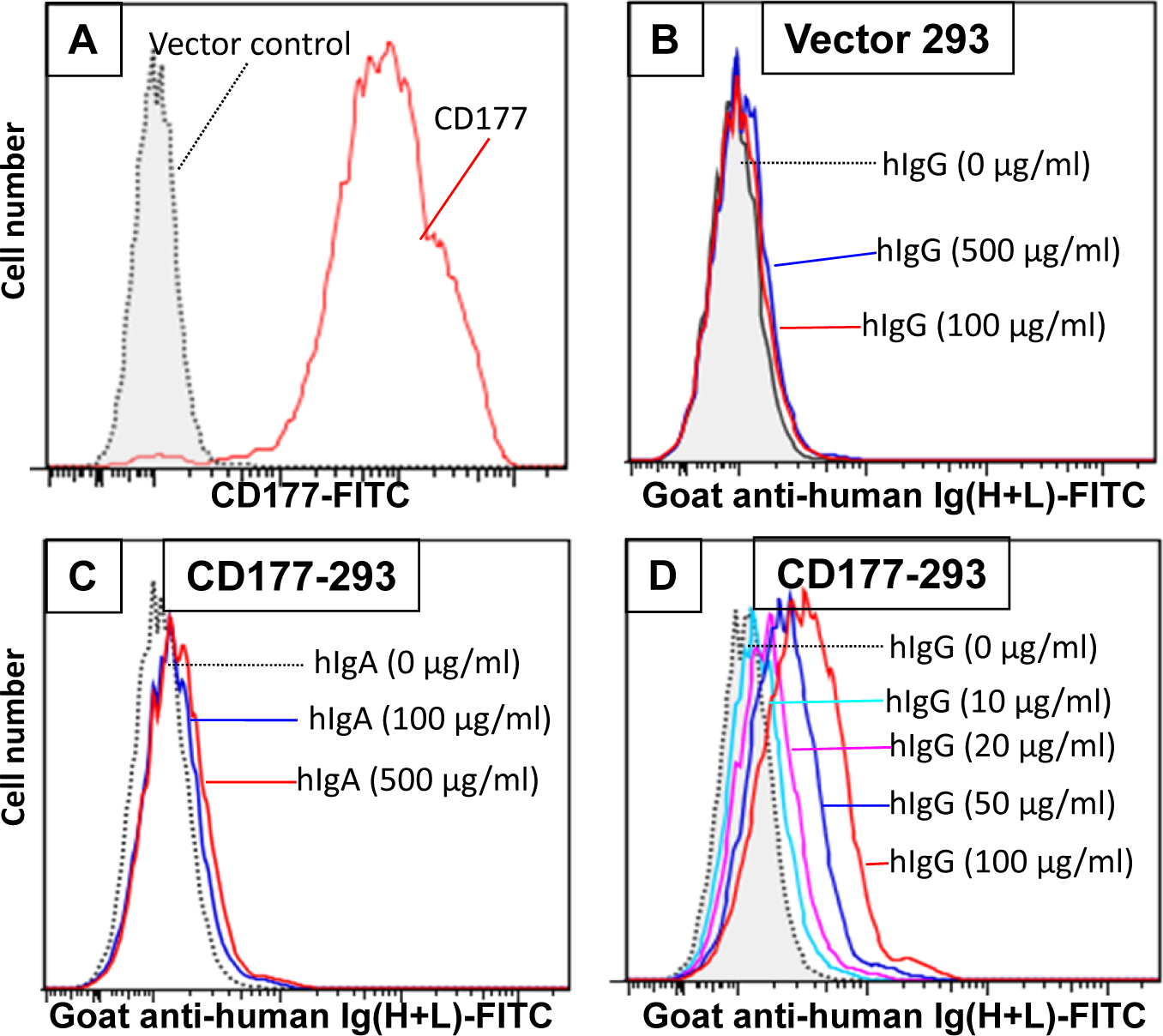
293 cells expressing CD177 bind human IgG (hIgG) in a dose-dependent manner. **A**. Expression of CD177 on 293 cells. 293 cells transfected with the vector do not express CD177. **B**. Vector control 293 cells were unable to bind hIgG. IgG binding assay was carried out as described in “Matrials and Methods”. No binding could be observed with hIgG at concentrations of 100 or 500 µg/ml. **C**. 293 cells expressing CD177 were unable to bind human IgA (hIgA). **D**. 293 cells expressing CD177 bound hIgG in dose-dependent manner. Increased hIgG concentrations led to more IgG bound on 293 cells expressing CD177.

**Fig. 2.**
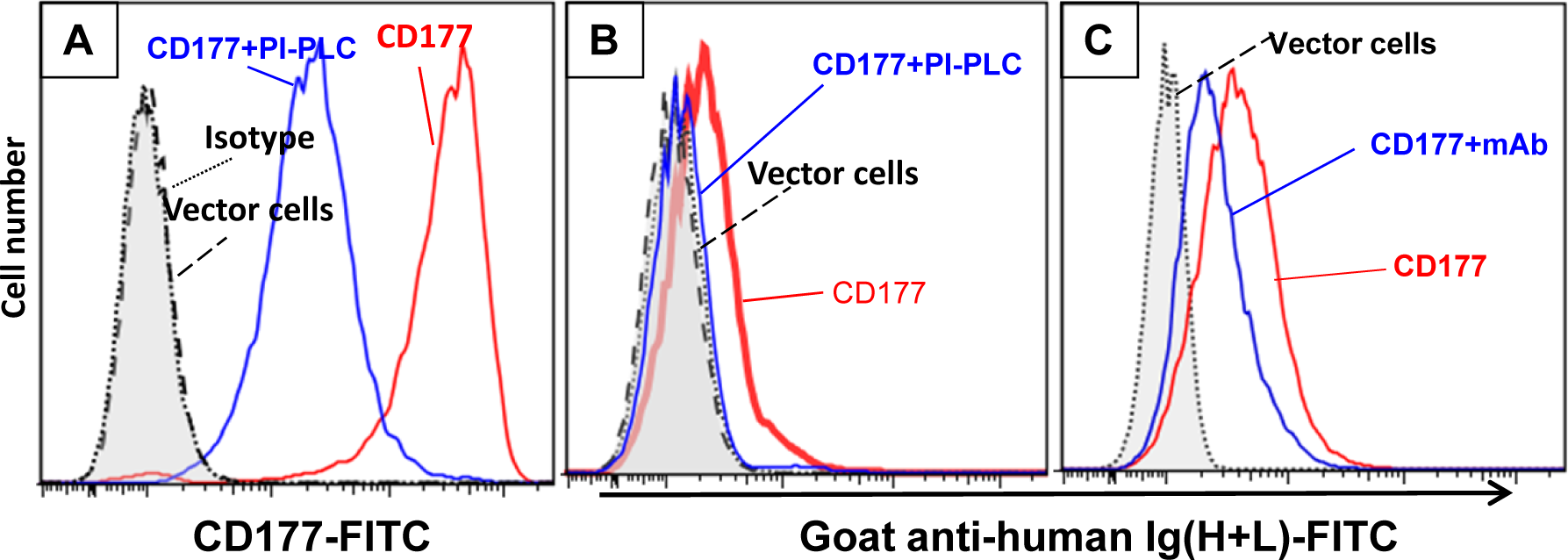
Cleavage of CD177 from cell surface and treatment of cells with anti-CD177 monoclonal antibody inhibit the binding of CD177 to hIgG. **A**. Treatment of the 293 cells expressing CD177 with PI-PLC (CD177+PI-PLC) removed most of CD177 from cell surface compared to the same cells without PI-PLC treatment (CD177). **B**. IgG binding assay was carried out as described in “Matrials and Methods”. The removal of CD177 from the cell surface (CD177+PI-PLC) abolished hIgG binding to cells compared to the same cells without PI-PLC treatment (CD177). **C**. Cells treated with anti-CD177 mAb (CD177+mAB) significantly inhibited hIgG binding compared to the cells without mAb treatment (CD177).

### CD177 is a low affinity IgG Fc receptor that bind all human IgG subclasses with similar affinity

To measure the affinity of CD177 for human IgG subclasses, we produced soluble CD177 (sCD177) containing CD177 extracellular domain (residue 1 – 408) (**Supplemental Fig. S1**) for surface plasmon resonance (SPR) analysis. In multicycle SPR kinetic analysis, different concentrations (125, 250, 500, 1,000, 2,000, 4,000, 8,000, and 16,000 nM) of sCD177 were injected into flow cells at a flow rate of 30 µl/min, with a contact and dissociation time of 250 and 600 seconds, respectively (**Fig. 3**). As shown in **Table 1**, all human IgG subclasses bound to CD177 ectodomains with a narrow range of affinity (kinetic K_D_ between 1.97×10^-6^ and 2.798×10^-6^ M and steady state K_D_ between 2.97×10^-5^ and 4.12×10^-5^ M). Results from the SPR assay demonstrate that CD177 is a low affinity IgG Fc receptor capable of binding to all human IgG subclasses.

**Fig. 3.**
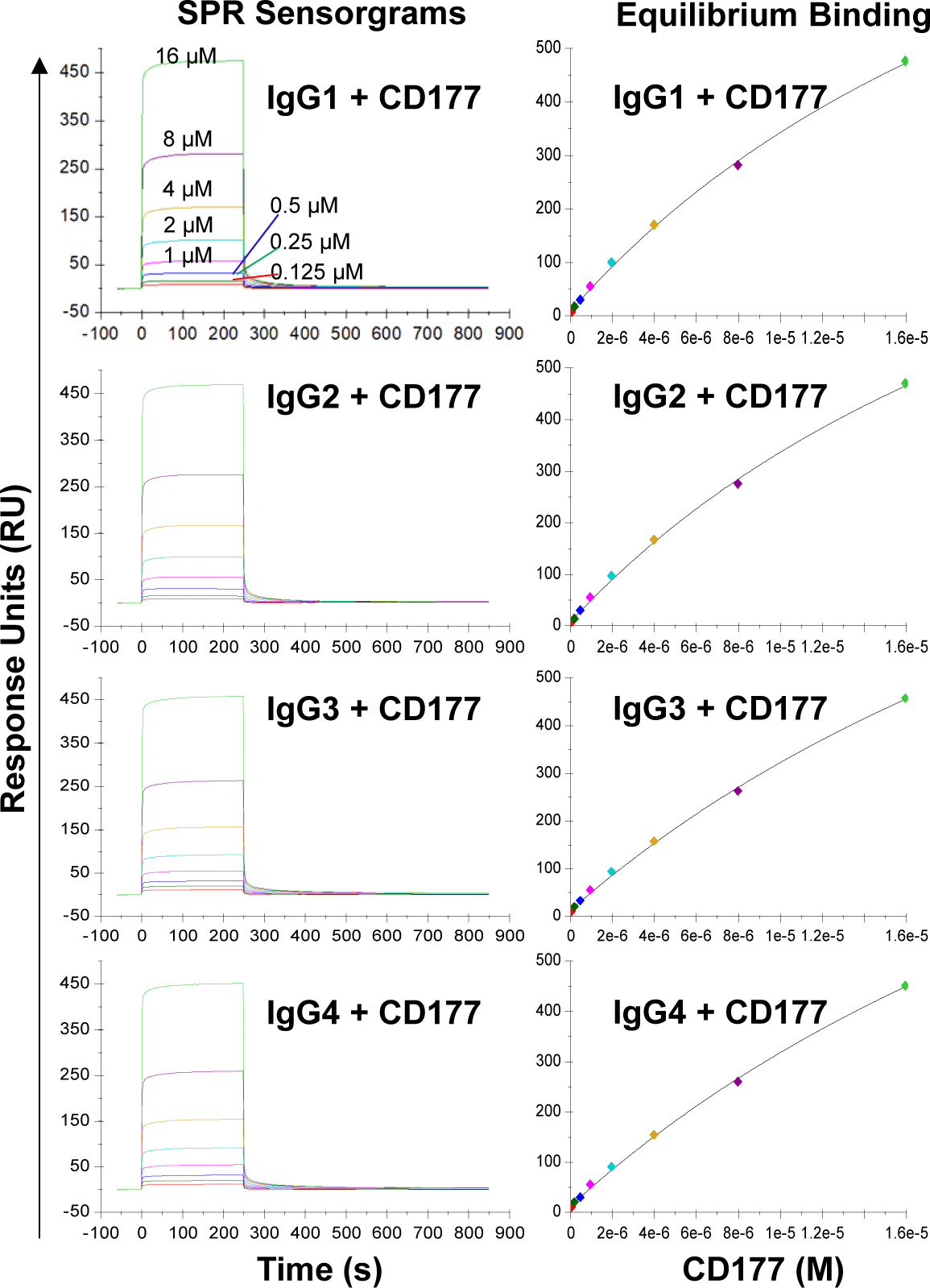
Binding of IgG subclasses to human CD177 by SPR. The left column shows representative SPR sensorgrams and the right column shows steady state binding curves. The experiments were repeated three times and equivalent results were obtained.

**Table 1.** Summary of affinity constants determined by equilibrium and kinetic analysis.

| **Ligand** | **Analyte** | **kon (1/Ms)** | **koff (1/s)** | **Kinetic K<sub>D</sub> (M)** | **Steady state K<sub>A</sub> (M)** |
| --- | --- | --- | --- | --- | --- |
| IgG1 | CD177 | 3042 | 0.005395 | 1.773E-6 | 2.97e-05 |
| IgG2 | CD177 | 3350 | 0.009372 | 2.798E-6 | 3.03e-05 |
| IgG3 | CD177 | 3070 | 0.004940 | 1.609E-6 | 4.07e-05 |
| IgG4 | CD177 | 3346 | 0.004006 | 1.197E-6 | 4.12e-05 |

### CD177 alleles affect receptor expressions and interactions with human IgG

We previously demonstrated that the common nonsense CD177 SNP 263K>* (rs201821720) is the primary genetic determinant for CD177 null and expression variations (22, 23). We also found that the CD177 SNP 431G>R (rs78718189) leads to low CD177 expression on neutrophils (23). Haplotype analyses revealed that the nonsense SNP 263* allele is in complete linkage disequilibrium with SNP 431G allele while 263K is in complete linkage disequilibrium with431R. These two CD177 SNPs form three haplotypes/alleles (263K431G, 263*431G, and 263K431R) (**Fig. 4**). All homozygous 263*431G genotype donors were CD177 deficient (**Supplemental Fig. 2**). In addition, homozygous 263K431R genotype donors (**Supplemental Fig. 2B**) had significantly lower CD177 density on neutrophils (**Supplemental 2D)** than homozygous 263K431G donors (**Supplemental Fig. 2C)**. We hypothesize that CD177 263K431R allele leads to low CD177 expression, which may affect the interaction between human IgG and CD177. To determine effect of CD177 alleles on the binding capacity to human IgG, we used 293 cells stably expressing three CD177 alleles and vector control (**Fig. 4A**) for IgG binding assay. The nonsense CD177-263*431G allele is unable to express CD177 on cell surface. On the hand, CD177-263K431R allele expressed a much lower level of CD177 on cell surface than the wild type CD177-263K431G allele (**Fig. 4A)**, consistent with CD177 phenotypes on neutrophils (**Supplemental 2D)**. Our data confirm that CD177 alleles significantly affect cell surface CD177 expressions and that CD177 263K431R is defective allele for CD177 expression as previously reported (23).

**Fig. 4.**
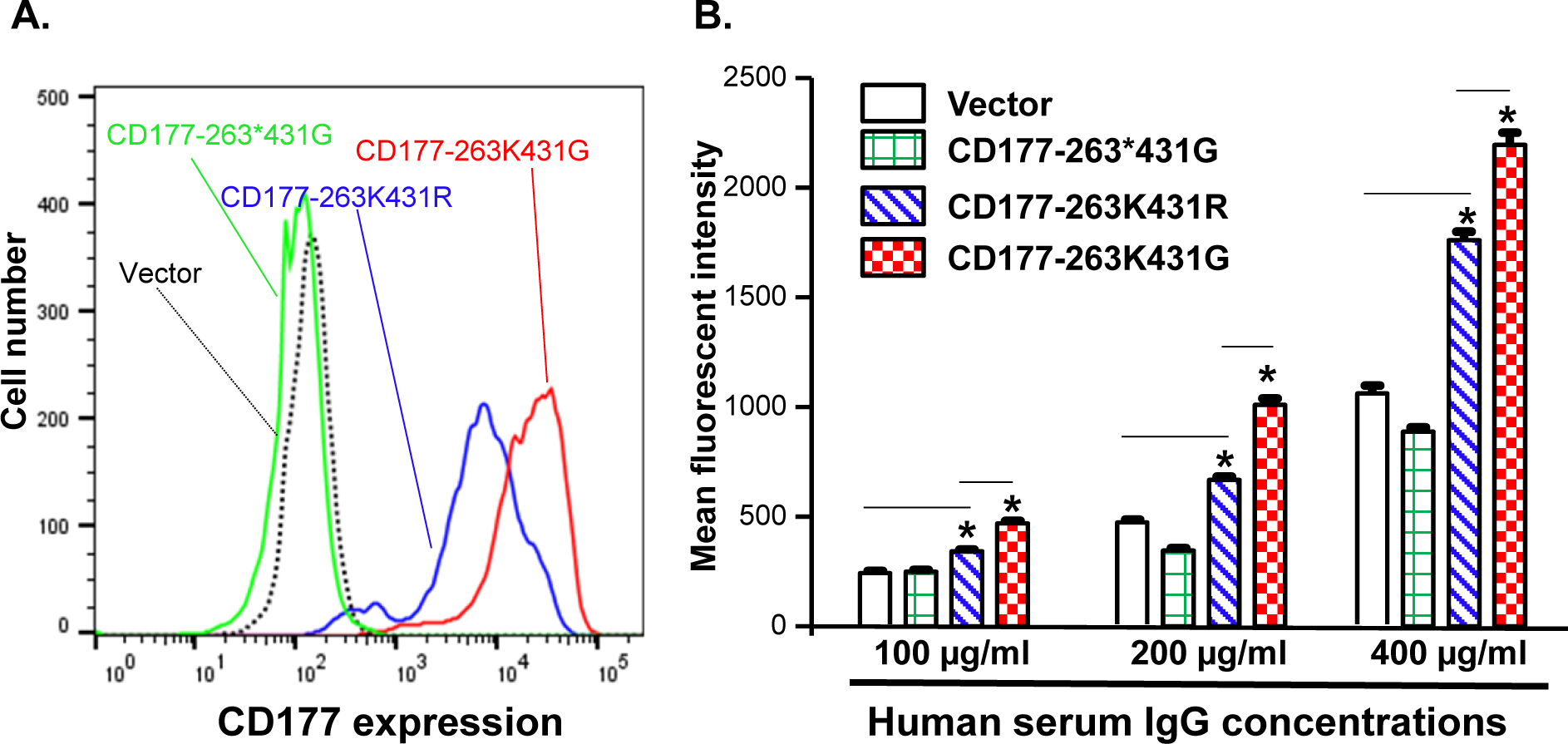
CD177 alleles affect CD177 expressions and binding to human IgG. **A**. 293 cell lines stably expressing three CD177 alleles (263K431G, 263*431G, and 263K431R) and vector control. CD177-263*431G cells do not express CD177 as compared to vector control cells. CD177-263K431R cells expressed much lower CD177 compared to CD177-263K431G cells. **B**. Binding capacity of cells with three CD177 alleles to hIgG. Flow cytometry-based IgG binding assay were carried out as described in “Materials and Methods”. The mean fluorescent intensities (MFI) represented the binding capacity of CD177 alleles for human IgG. CD177-263*431G cells failed to express CD177 on 293 cells were unable to bind hIgG, similar to vector control cells. CD177-263K431R cells had significantly lower binding capacity compared to CD177-263K431G cells. Results Data were means ± SD of three independent experiments (* *P* < 0.05).

Subsequently, we used flow cytometry-based binding assays to determine IgG binding capacity of 293 cell lines stably expressing CD177 alleles (263K431G, 263*431G, and 263K431R). As shown in **Fig. 4B**, the common 263K431G allele had the highest IgG binding capacity among three CD177 alleles. Cells expressing nonsense CD177-263*431G allele were unable to bind IgG as mean fluorescent intensities of those cells were statistically not different from those of vector control cells under various IgG concentrations. On the other hand, cells expressing 263K431R allele had significantly lower binding capacity than those expressing the wild-type 263K431G allele for IgG at the concentrations of 100, 200, and 400 µg/ml. Our data demonstrate that CD177 alleles significantly affect cell surface CD177 expressions and the interaction with human IgG.

### CD177 alleles affect binding capacity to human IgG-coated particles

We carried out beads-rosette formation assay to determine the ability of CD177 to bind IgG-coated beads (particle targets) and effects of CD177 alleles on IgG binding. As shown in **Fig. 5D**, cells expressing the wild-type CD177-263K431G allele are capable of forming typical beads-rosettes with multiple IgG-bound beads on a single cell (≥ 4 beads per cell). On the other hand, large beads-rosettes (a single cell bound more than four IgG-coated beads) were absent with the cells expressing nonsense 263*431G allele (**Fig. 5B**), similar to the vector control cells (**Fig. 5A**). In addition, cells expressing CD177-263K431R allele also could not effectively form large rosettes with IgG-coated beads (**Fig. 5C**). To quantitate the binding capacity of three CD177 alleles for IgG-coated beads, bead number on cells with the attachment of two or more beads was counted from one hundred random cells. As shown in **Fig. 5E**, cells expressing CD177-263K431G bound significantly more IgG-coated beads (123 ± 12) than those expressing CD177-263*431G (15 ± 4) and CD177-263K431R (13 ± 5) cells alleles (*P* < 0.0001). A low number of beads also bound to vector control cells (10 ± 6) due to the background or non-specific interaction between beads and cells. Interestingly, no significant differences of bound beads per 100 cells were observed among vector control, nonsense CD177-263*431G, and CD177-263K431R cells (*P* > 0.05). Our data clearly demonstrate that CD177 is an IgG receptor and that CD177 alleles significantly affect the binding capacity of CD177 to IgG-bound particles.

**Fig. 5.**
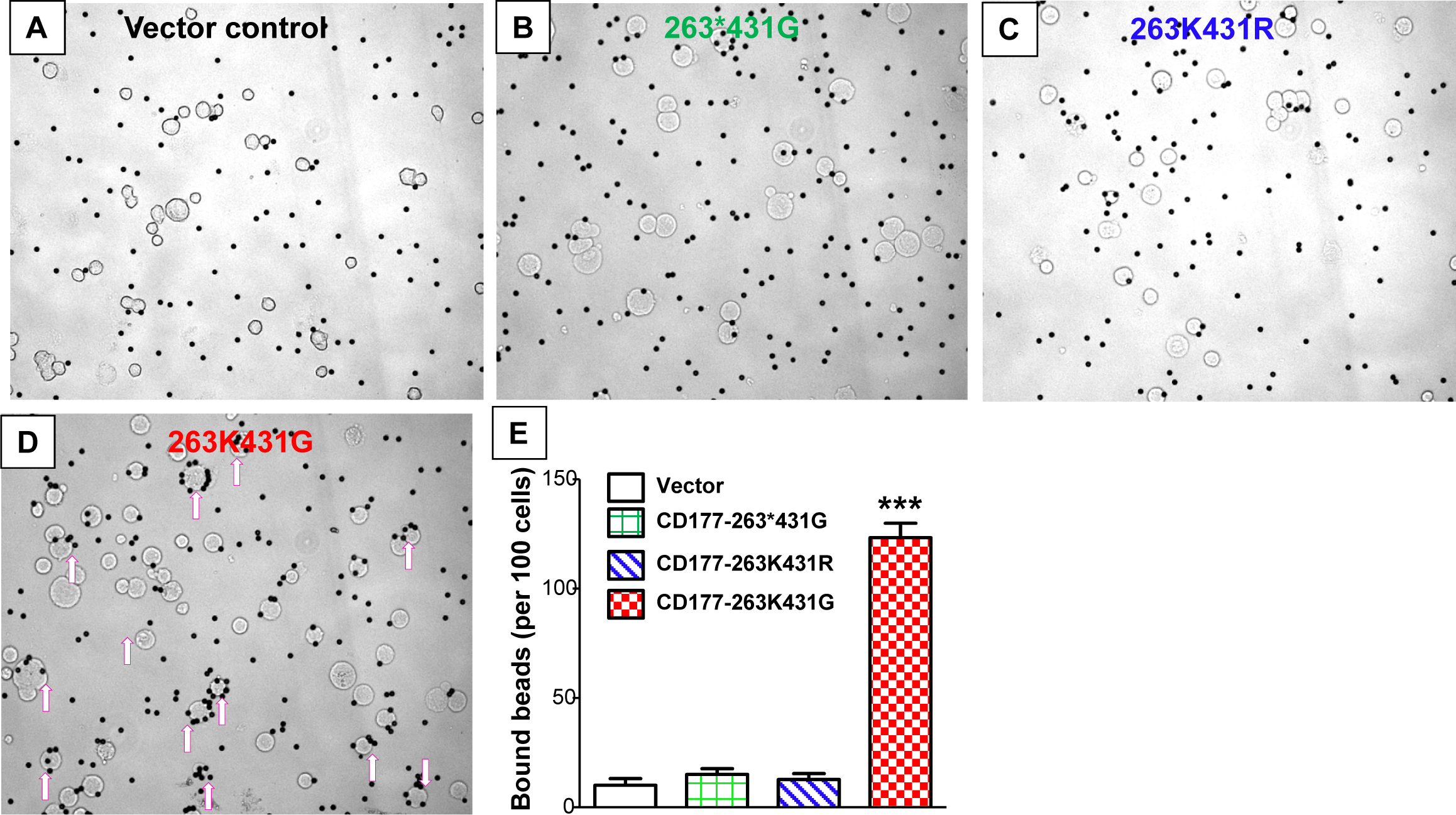
CD177 alleles affect binding capacity for IgG-coated particles. Beads-rosette formation assay was carried out to determine the ability of CD177 to bind IgG-coated particles. Large beads-rosettes (a single cell bound more than four IgG-coated beads) were absent for vector control cells (**A**), nonsense 263*431G allele cells (**B**), and 263K431R allele cells (**C**) while cells with 263K431G allele are capable of forming large beads-rosettes (arrows pointed cells) (**D**). **E**. Cells expressing 263K431G allele bound significantly more IgG-coated beads (123 ± 12) than those expressing CD177-263*431G (15 ± 4) and CD177-263K431R (13 ± 5) cells alleles (*P* < 0.0001). Results were mean ± SD of three experiments.

### CD177 alleles affects ADCC function

To determine effect of CD177 alleles on immune function, we generated three CD177-pBMN-IRES-EGFP retroviral constructs (263K431G, 263*431G, and 263K431R) and established NK-92 stable cell lines expressing CD177 alleles. CD177-263*431G allele failed to express CD177 on NK-92 cells and NK-92 cells with CD177-263K431R allele express a much lower density of CD177 compared to those with wild-type CD177-263K431G allele (**Fig. 6A**), consistent with the phenotypes of those two alleles in peripheral blood neutrophils (**Supplemental Fig. 2**). Both CD177-263K431R and nonsense 263*431G allele failed to mediate ADCC in NK-92 cells and the percentages of specific release (or ADCC activity) are near zero and similar to those without Trastuzumab. On the other hand, cells expressing the common CD177-263K431G allele had significantly higher ADCC activity (% specific release = 8.93 ± 4.30%) than those expressing either 263K431R (% specific release = 0.11 ± 0.29 or 263*431G alleles (% specific release = 0.04 ± 0.05) (**Fig. 6B**) (*P* = 0.0004). CD177-263K431G allele cells has a higher background natural cytotoxicity (% specific release = 1.45 ± 1.42) as compared to 263K431R (% specific release = 0.03 ± 2.31) or 263*431G alleles (% specific release = 0.03 ± 0.06) in the absence of Trastuzumab, but the differences were statistically significant (*P* = 0.2224). Our data suggest that CD177 is a functional IgG Fc receptor capable of mediating ADCC and CD177 alleles affect IgG-mediated function.

**Fig. 6.**
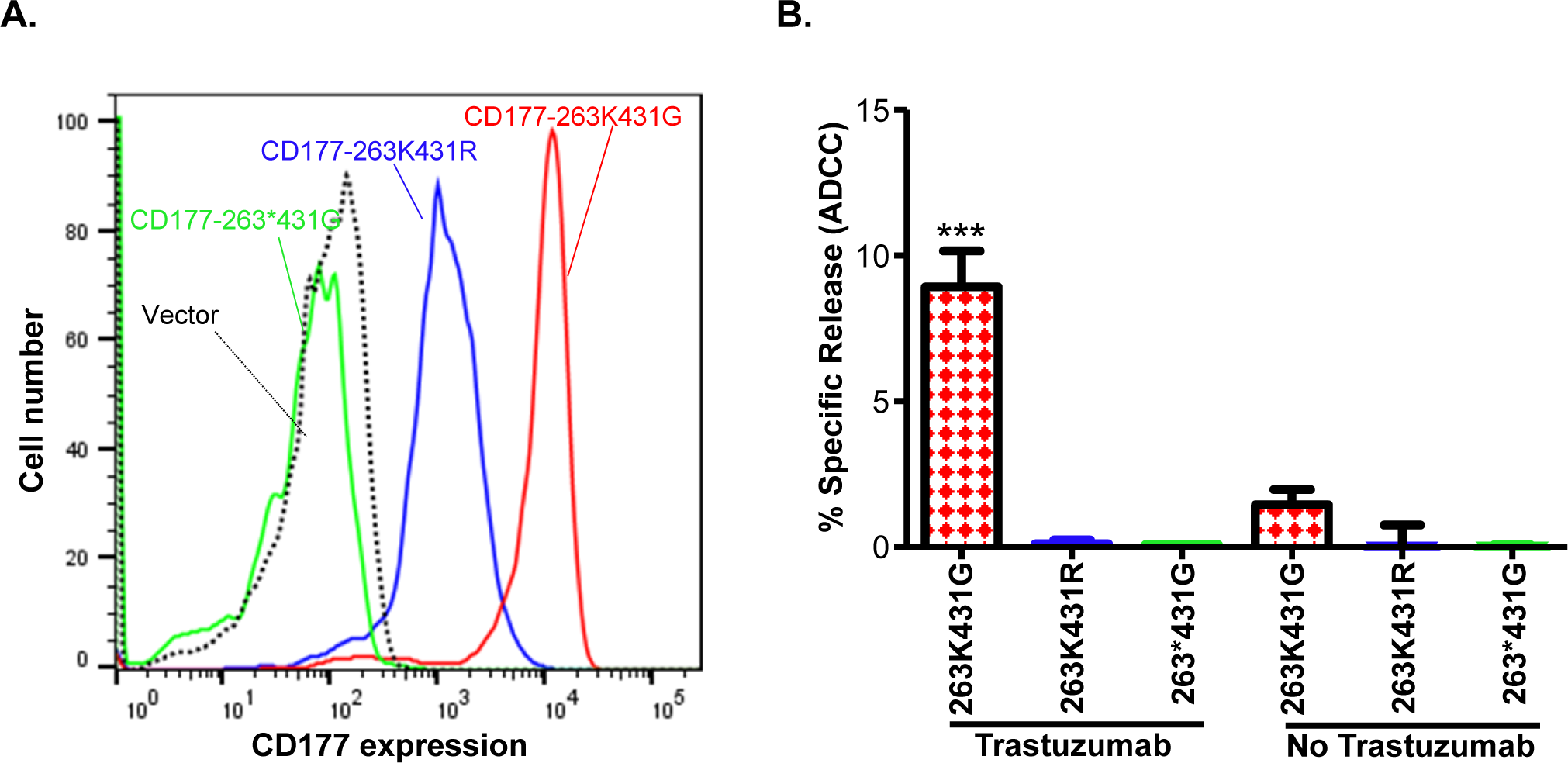
CD177 alleles affect surface CD177 expression and ADCC function of NK-92 cells. **A**. Expressions of CD177 on stably NK-92 cell lines carrying CD177 alleles. NK-92 cells carrying 263*431G allele failed to express CD177 on NK92 cells. NK-92 cells with 263K431R allele express a much lower density of CD177 compared to those with 263K431G allele. **B**. ADCC activity of NK-92 cells expressing CD177 alleles. NK-92 cells expressing 263K431G allele had the highest ADCC activity among three CD177 alleles (*P* < 0.001). Both 263K431R and the null allele CD177-263*431G failed to mediate ADCC, similar to the vector control cells. Results from four experiments are shown.

## Discussion

Inherited CD177 deficiency is common in human populations but its potential impact on immune functions remains unknown. In the current study, we revealed CD177 as a novel IgG Fc receptor that mediates ADCC function. Notably, we observed that CD177 genetic alleles significantly affect its interaction with human IgG and ADCC functions. The definition of CD177 as novel functional IgG Fc-receptor may fill a knowledge gap about the full repertoire of human FcγRs. Our study provided novel insights into CD177-mediated immune functions and into the potential genetic mechanisms underlying variability of responses to vaccination and antibody therapy.

Our experimental data show that CD177 is a low affinity IgG receptor. However, CD177 differs from classical low affinity IgG Fc receptors in three features. First, the binding affinity of CD177 for human IgGs (steady state K_D_ between 2.97×10^-5^ and 4.12×10^-5^ M) is much lower than those of the classical low-affinity IgG Fc receptors (the steady state K_D_ between 10^-6^ to 10^-7^) (30). Second, CD177 binds all IgG subclasses with similar affinity while classical IgG Fc receptors bind IgG subclasses with significantly different affinities (30). Third, the reduction of CD177 density on cell surface had a dramatic effect on the interaction between CD177 and IgGs, which could be explained by very low affinity of CD177 for IgGs compared to classical low affinity FcγRs.

The common nonsense CD177 SNP 263K>* is the primary genetic determinant for CD177 null and expression variations (37) while the CD177 SNP 431G>R also leads to low CD177 expression on neutrophils (23). The nonsense SNP 263* allele is in complete linkage disequilibrium with SNP 431G allele and 263K is in complete linkage disequilibrium with 431R, leading to three haplotypes or alleles (263K431G, 263*431G, and 263K431R in human population. All homozygous 263*431G genotype donors were CD177 deficient (37). In addition, homozygous 263K431R genotype donors had significantly lower CD177 density on neutrophils than homozygous 263K431G donors (23). In the current study, CD177-263K431R allele consistently leads to lower CD177 expression than the common/wild-type CD177-263K431G allele in stable transfected 293 and NK-92 cell lines, confirming that CD177-263K431R is also a defective CD177 allele for expression. Importantly, cells expressing 263K431R allele had significantly lower binding capacity for human IgG and IgG-coated beads compared to those expressing the 263K431G allele due to low expression of 263K431R allele on cell surface. Notably, CD177-263K431R allele failed to mediate ADCC, indicating that CD177-263K431R allele not only affects CD177 surface expression and binding capacity for IgG but also influence the CD177 immune function. We conclude that both CD177 SNP 263K>* and SNP 431G>R affect CD177 functions.

The function of CD177 remains poorly understood. CD177 is a GPI-anchored membrane protein tethered to the neutrophil surface by a glycosylphosphatidylinositol anchor lacking a transmembrane domain. CD177 employs Mac-1 (CD11b/CD18) to transduce signals for cell functions (24, 33–35), reminiscent of the low affinity IgG Fc receptor FcγRIIIB (36–39). CD177 is selectively expressed in bone marrow cells and plays an important role in the proliferation and differentiation of myeloid lineage cells including neutrophils, myelocytes, promyelocytes, megakaryocytes, and early erythroblasts. In a CD177-genetic knockout mouse model, CD177-deficiency leads to increased neutrophil cell death in bone marrow, suggesting that CD177 could provide survival signal during neutrophil differentiation (13). CD177 is over-expressed in neutrophils from patients with myeloproliferative disorders including polycythemia vera, essential thromobocythemia, idiopathic myelofibrocythemia, and hypereosinophilic syndrome (14, 18). CD177 was an indicator of increased erythropoietic activity in thalassemia syndromes as CD177 expression was significantly elevated in β-thalassemia patients compared to healthy controls (40). CD177 overexpression may also have a direct role in the pathogenesis of myeloproliferative disorders as CD177 enhances cell proliferation in vitro (11, 12). On the other hand, the low percentage of CD177 positive neutrophil is significantly associated with myelodysplastic syndrome and chronic myelogenous leukemia (10, 17). In vitro studies show that CD177 enhances cell growth/proliferation (11, 12). The current study suggests that IgGs may be natural ligands for CD177. As a low affinity IgG Fc receptor (FcγR), CD177 may only recognize multivalent IgG complexes. Accordingly, CD177 could serve as a sensor for IgG immune complexes (or IgG-opsonized targets) in bone marrow or in circulation. We speculate that that interaction between CD177 of myeloid progenitor cells and IgG immune complexes may stimulatory signal to drive promyeloid cell proliferation and differentiation into mature neutrophils during infections and diseases. Nevertheless, the exact role of interaction between CD177 and IgG immune complexed in myeloid cell development requires further investigation. Future studies also need to focus on direct effects of CD177 expression and deficiency on immune system in human subjects.

### Conclusion

CD177 is a novel IgG Fc receptor and the CD177 genetic variants significantly affect the binding of IgG. Additionally, CD177 could mediate ADCC function and genetic variants of CD177 significantly affect the immune function. The definition of CD177 as novel functional IgG Fc-receptor fills the critical knowledge gap about the full repertoire of human FcγRs. Our study may provide novel insights into CD177-mediated immune functions and into the genetic mechanisms underlying response variability of antibody therapy. CD177 genetic variants may also serve as valuable biomarkers in antibody therapy and vaccinations.

## AUTHOR CONTRIBUTIONS

Both authors contributed to the study conception and design. Conceptualization, resources, funding acquisition: Jianming Wu. Methodology, investigation, data acquisition and analyses: Jianming Wu, Yunfang Li. Manuscript preparation: Jianming Wu. All authors have read and agreed to the published version of the manuscript.

## CONFLICT OF INTEREST

The authors have no relevant financial or non-financial interests to disclose.

**ORCID:** Jianming Wu – http://orcid.org/0000-0001-9142-7066.

---

**Supplementary Fig. S1.**
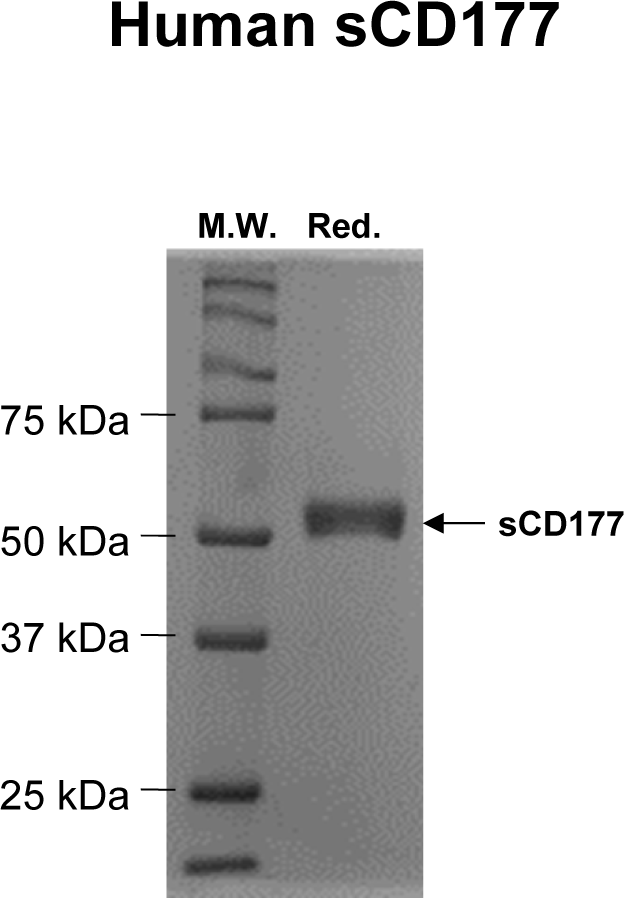

**Supplementary Fig. S2.**
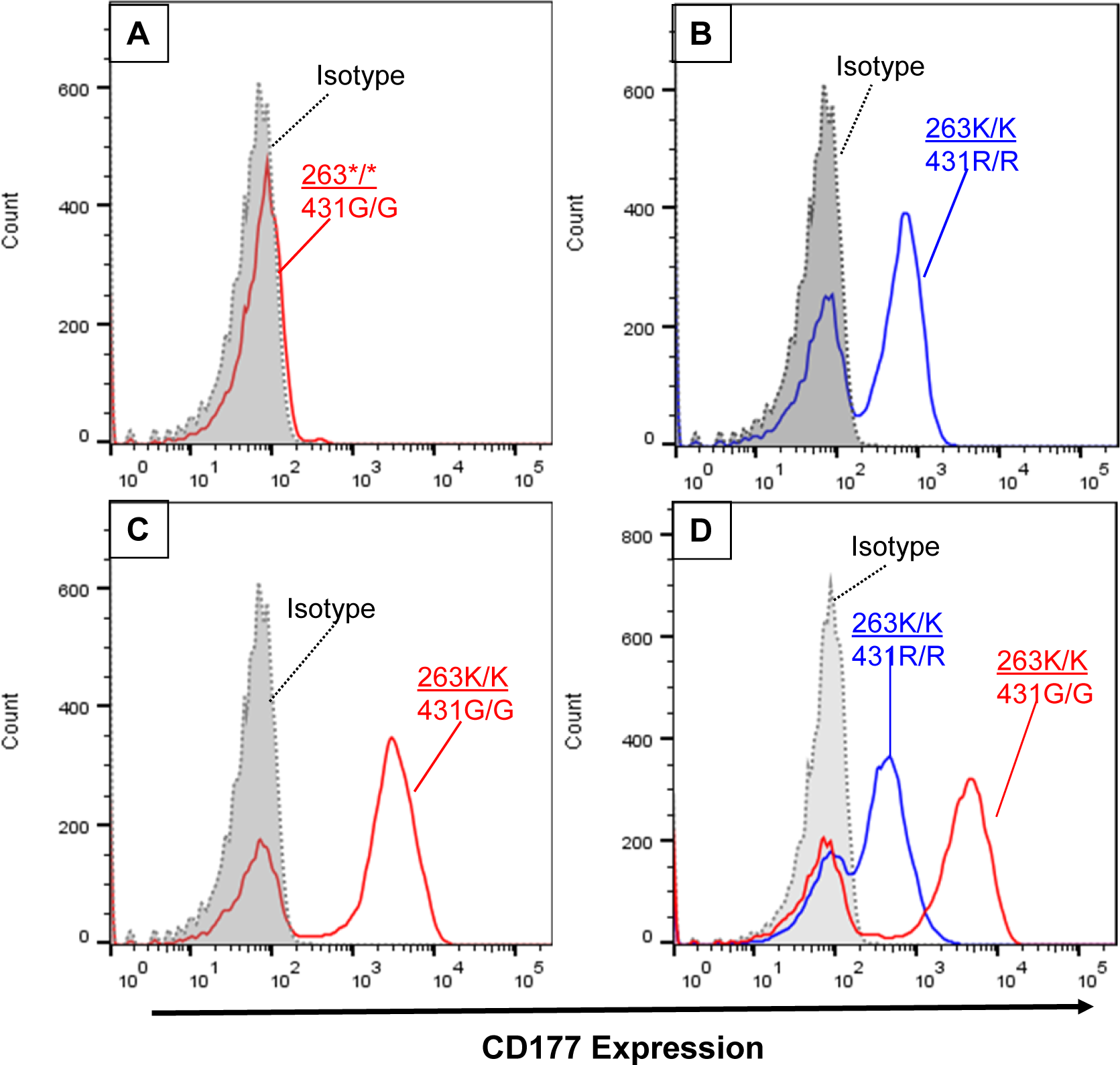

